# γ-proteobacteria eject their polar flagella under nutrient depletion, retaining flagellar motor relic structures

**DOI:** 10.1101/367458

**Authors:** Josie L. Ferreira, Forson Z. Gao, Florian M. Rossmann, Andrea Nans, Susanne Brenzinger, Rohola Hosseini, Ariane Briegel, Kai M. Thormann, Peter B. Rosenthal, Morgan Beeby

## Abstract

Bacteria switch only intermittently to motile planktonic lifestyles under favourable conditions. Under chronic nutrient deprivation, however, bacteria orchestrate a switch to stationary phase, conserving energy by altering metabolism and stopping motility. About two-thirds of bacteria use flagella to swim, but how bacteria deactivate this large-molecular machine remains poorly studied. Here we describe the previously unreported ejection of polar sodium-driven motors by γ-proteobacteria. We show that these bacteria eject their flagella at the base of the flagellar hook when nutrients are depleted, leaving a relic of a former flagellar motor in the outer membrane. Subtomogram averages of the full motor and relic reveal that this is an active process, as a plug protein appears in the relic, likely to prevent leakage across their outer membrane. We show that this is a widespread phenomenon demonstrated by the appearance of relic structures in varied γ-proteobacteria including *Plesiomonas shigelloides, Vibrio cholerae, Vibrio fischeri, Shewanella putrefaciens* and *Pseudomonas aeruginosa*.

## Introduction

Many bacteria switch to a non-motile lifestyle in stationary phase to conserve energy (Adler and Templeton, 1967), but how this switch is accomplished is poorly understood. The most widespread motility device used by bacteria is the flagellum, found in approximately two thirds of bacteria (Chen et al., 2011). Bacterial flagellar filaments are helical propellers that extend multiple microns from the cell from a periplasm-spanning rotary motor. Motor torque is generated by harnessing ion flux across the inner membrane; torque is transmitted to a periplasm-spanning rod to the extracellular filament. The rod exits the cell through dedicated P- and L-rings that act as channels through the peptidoglycan layer and outer membrane, respectively. When rotated, the filament acts as a propeller that exerts thrust for propulsion of the bacterium. Although the flagellum is vital for migration to favourable environments, sites of biofilm formation, or sites of infection, however, it is counterproductive for the cell to retain a functional flagellum during nutrient depletion, and stopping motility is preferable to resource exhaustion.

Methods for deactivation of flagella in environments other than nutrient depletion are diverse across those bacteria studied to date. *Rhodobacter sphaeroides* has a unidirectional flagellum that is stopped by a “molecular brake” for navigation (Pilizota et al., 2009) while *Bacillus subtilis* uses a “molecular clutch” to stop flagellum rotation and swimming for biofilm formation (Blair et al., 2008). The α-proteobacterium *Caulobacter crescentus*, meanwhile, actively ejects its single, polar flagellum upon surface sensing (Ellison et al., 2017; Hug et al., 2017) in order to build an adhesive stalk for surface adhesion.

Little is currently known about how the widespread polar-flagellated γ-proteobacteria modulate motility. This group includes a diverse set of pathogens occupying both sessile and planktonic niches, including human pathogens *Vibrio cholerae* and *Plesiomonas shigelloides*, opportunistic pathogens *Pseudomonas aeruginosa* and *Shewanella putrefaciens*, and non-pathogenic members including the squid symbiont *Vibrio fischeri*. These γ-proteobacteria have one or more polar flagella powered by sodium ions that are faster (Magariyama et al., 1994) and higher-torque (Beeby et al., 2016) than the model flagellar motor from *Salmonella*. Sodium-driven polar motors have a number of structural differences to the well-studied enteric-like motors (Beeby et al., 2016; Terashima et al., 2013; Zhu et al., 2017a). Most striking in subtomogram averages are the addition of periplasmic (H-ring and T-ring) and outer membrane (basal) disks (Beeby et al., 2016; Chen et al., 2011). The T-ring, made up of MotX and MotY contributes to the assembly and scaffolding of the stator complexes (PomA/PomB). Many γ-proteobacteria also incorporate a sheath, an outer membrane-extension that encapsulates the flagellum; the H-ring and basal disk, composed of FlgO, FlgP, and FlgT, has recently been shown to assist in outer membrane penetration in these bacteria (Liu et al., 2018), although its function is likely broader than sheath formation given that many unsheathed bacteria also retain an H-ring.

Here, we describe work showing that γ-proteobacteria eject their sodium-driven polar flagella in response to nutrient depletion. The ejected flagella leave a “relic” of the ejected motor in the outer membrane composed of the P-, L-, H- and T-rings and the basal disk. During this transition a previously undescribed plug is incorporated into the relic, likely to block periplasmic leakage, and suggesting an active, controlled mechanism. We speculate on the nature of the novel flagellar plug and the significance of flagellar ejection.

## Results

### γ-proteobacterial motility slows as cell density increases

While tracking the swimming speeds of γ-proteobacterium *Plesiomonas shigelloides*, we noticed that bacteria with multiple sodium-driven polar flagella showed a significant decrease in swimming speed at high cell densities (Fig. 1a). Video tracking of individual bacterial cells revealed that *P. shigelloides* swam at 40 μm s^−1^ between OD 0.2 and OD ^∼^0.7 before swimming speeds sharply dropped at OD 0.8, down to 12 μm s^−1^ at OD 1.0. Furthermore, the percentage of active swimmers dropped from over 95% at low OD values to approximately 5% by OD 1.0. Another γ-proteobacterium, *Vibrio fischeri*, displayed the same characteristic behavior (Fig. 1a). Strikingly, however, the model enteric bacterium *Salmonella enterica* subsp. enterica serovar Typhimurium (*Salmonella)* that uses a different family of flagellar motors continued swimming as well as, if not faster than, *Salmonella* cells at OD 0.2 when cultured to higher cell densities (Fig. 1a).

**Figure 1:**
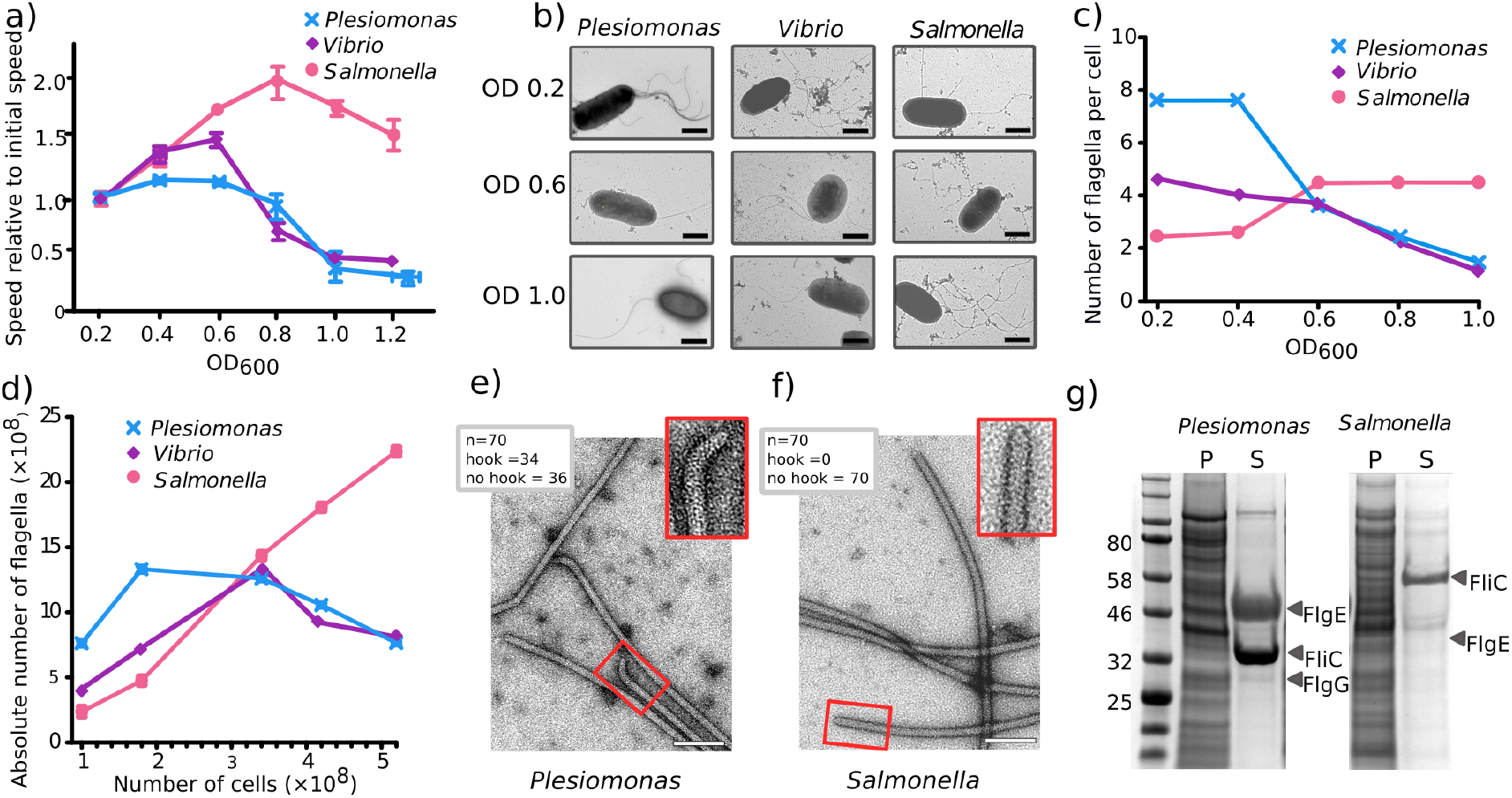
γ-proteobacteria swimming slows at high cell densities due to loss of flagella. a) Swimming speeds of *Plesiomonas shigelloides, Vibrio fischeri* and *Salmonella enterica* sv Typhimurium at increasing cell density. Speed relative to initial speed at OD_600_ 0.2 are represented. Error bars indicate standard error. b) Representative negative stain EM images of *P. shigelloides, V. fischeri* and *S. enterica* at three different cell densities. Scale bars are 1µm. c) Mean number of flagella, counted from 150 cells (50 per biological replicate) at increasing cell densities. d) The absolute number of flagella in the population calculated from the mean number of flagella and CFU. Error bars indicate standard error. e) Negative stain EM images of flagella recovered from the supernatant of stationary phase *P. shigelloides*. Inset shows close-up of a hook. Scale bar is 100nm. f) Negative stain EM images of flagella recovered from the supernatant of stationary phase *S. enterica*. Inset shows close-up of a broken filament. Scale bar is 100nm. g) SDS-PAGE gel of *P. shigelloides* and *S. enterica* cells grown to stationary phase. Polar FliC (filament) and FlgE (hook) proteins are seen in the supernatant of the P. shigelloides culture. S = supernatant, P = pellet.

### Decreased swimming speed is due to loss of flagella

In the course of a previous study (Beeby et al., 2016) we noted that flagellation levels in γ-proteobacteria are highest at low cell optical densities. To determine whether the decrease in swimming speed of γ-proteobacteria was due to differences in flagellation levels at different cell densities, flagella were counted in negative stain EM images of cells to assess flagellation levels (Fig. 1b). Time-courses of both *P. shigelloides* and *V. fischeri* cells at increasing cell-densities showed decreasing numbers of flagella per cell (Figure 1b and c) and cells grown overnight into late stationary phase had no flagella. *Salmonella*, in contrast, had increasing or stable numbers of flagella at higher cell densities.

We sought to distinguish whether solely flagella synthesis is down-regulated, or whether flagella are also actively disassembled in γ-proteobacteria at high cell densities. Plotting the absolute number of flagella in the entire population suggested that flagella are being ejected. The number of cells in a population was determined by calculating colony forming units (CFU) and relating this to the mean number of flagella per cell as counted by EM (Fig. 1d). The absolute number of flagella in the *P. shigelloides* and *V. fischeri* populations declined overtime, demonstrating that bacteria are losing flagella faster than they are synthesizing them (Fig. 1d). The *Salmonella* positive control, in contrast, shows an increase in the absolute number of flagella in the population in direct correlation with CFU count, as assembly continues and no ejection occurs at higher cell densities.

### Used γ-proteobacterial growth medium contains free flagella with hooks at one end

To confirm that flagella are being ejected, we determined whether supernatant of *P. shigelloides* cells grown to stationary phase contains released flagella. PEG precipitation revealed that *P. shigelloides* growth medium indeed contained free flagella (Fig. 1e). Curiously, 50% of all flagella ends had an attached hook, leading us to infer – given that hooks are only present at the proximal end of all flagella, i.e., 50% of ends – that effectively all isolated *P. shigelloides* flagella have attached hooks, consistent with ejected flagella from *Caulobacter crescentus* (Kanbe et al., 2005). Flagellar filaments were also recovered from the supernatant of a *Salmonella* control (Fig. 1f), but no hooks were observed of 70 randomly imaged *Salmonella* filament ends.

### Shed hook-filament structures accumulate in *P. shigelloides* supernatant

To verify that *Salmonella* filaments are sheared mid-filament but *P. shigelloides* filaments are cleaved at the base of the hook, flagella were recovered from cell cultures and analysed by SDS-PAGE. Cells grown to high OD were removed by pelleting, and flagella recovered from the remaining supernatant by PEG precipitation before SDS-PAGE analysis. Similar samples were collected from *Salmonella* cultures and volumes adjusted to match *P. shigelloides* cell count. The *Plesiomonas* supernatant had significant bands at molecular weights corresponding to both the polar flagellin (FliC) and hook (FlgE), and a very faint band at the expected size of the distal rod (FlgG). The *Salmonella* supernatant contained a weaker flagellin band, and no band at the molecular weight corresponding to the *Salmonella* hook (FlgE) (Fig. 1g). This confirms that *Plesiomonas* cells consistently lose their flagella at the base of the hook, while *Salmonella* flagella accumulation in the supernatant is due to shearing mid-filament. We conclude that this consistent behavior in response to a specific environment, i.e., high cell densities, is best explained by an ejection mechanism.

### Apparent flagellar relics remain at the cell pole at high OD

The hypothesis that γ-proteobacteria eject their flagella led us to speculate that a partial flagellar motor structure would remain after ejection of the filament and hook. Cryo-electron tomograms were collected of *P. shigelloides* cells grown to either low or high cell density and inspected. Consistent with our hypothesis, partial flagellar structures were seen to remain at the pole of cells from high OD cultures with fewer in cells from low OD cultures. Our swimming speed and flagellar ejection results lead us to conclude that these structures are likely to be the “relics” of once fully-functioning flagella that were subsequently ejected in high OD cultures, leaving only the outer membrane rings (the P-, L-, H- and T-rings and the basal disk) assembled (substantiated further by a subtomogram average structure described below). The P- and L-rings, which are integral parts of the relic structure, have been shown to require a rod and chaperone for assembly, incapable of assembling in the absence of an assembled rod (Kubori et al., 1992).

The two datasets demonstrated that relics are enriched at high OD and therefore correlate with flagellar ejection (Table 1). When cells were cryo-preserved at OD 0.25, 43% of cell poles had exclusively intact flagella while 48% had both intact flagella and relics, and 9% had no structures. The average number of full flagella per pole was 5.5 and, if the cells had relics, the average number was 2. When cells were frozen at OD 1.0, however, none of the cells had exclusively intact flagella, 36% had both relics and intact flagella, 12% had exclusively relics, and 52% had none of either structure. In contrast to cells grown at low OD, if the cells had intact flagella the average seen was 2; the average number of relics was 5. These results link OD to presence of the relic structures and demonstrate that not only are flagella being ejected, but assembly is also stopping, as many cells at high OD have neither flagella nor relics, indicating newly formed poles that have not yet assembled a single flagellum.

**Table 1:** Appearance of outer membrane relic structures is related to cell density. Number of filaments and outer membrane relics counted from tomograms of *P. shigelloides* cells grown to OD_600_0.25 (N=44) or OD_600_1.0 (N=25).

| OD<sub>600</sub> | Cells with filaments and relics | Cells with only filaments | Cells with only relics | Cells with nothing | Average number of filaments | Average number of relics |
| --- | --- | --- | --- | --- | --- | --- |
| 0.25 | 48% | 43% | 0% | 9% | 5.5 | 2 |
| 1 | 36% | 0% | 12% | 52% | 2 | 5 |

### Diverse γ-proteobacteria retain relics of ejected flagella

In order to determine whether this pattern is widespread amongst the γ-proteobacteria beyond *P. shigelloides* we collected cryo-tomograms from four additional γ-proteobacterial species: *Vibrio cholerae, Vibrio fischeri, Shewanella putrefaciens* and *Pseudomonas aeruginosa*. Markedly, in all species, similar relic structures in the outer membrane of the poles was seen alongside intact flagella (Fig. 2a). This is particularly notable in *V. cholera* given that this species characteristically only ever has one flagellar motor at the pole. In one tomogram we observed four relics alongside an intact flagellum in *V. cholerae*.

**Figure 2:**
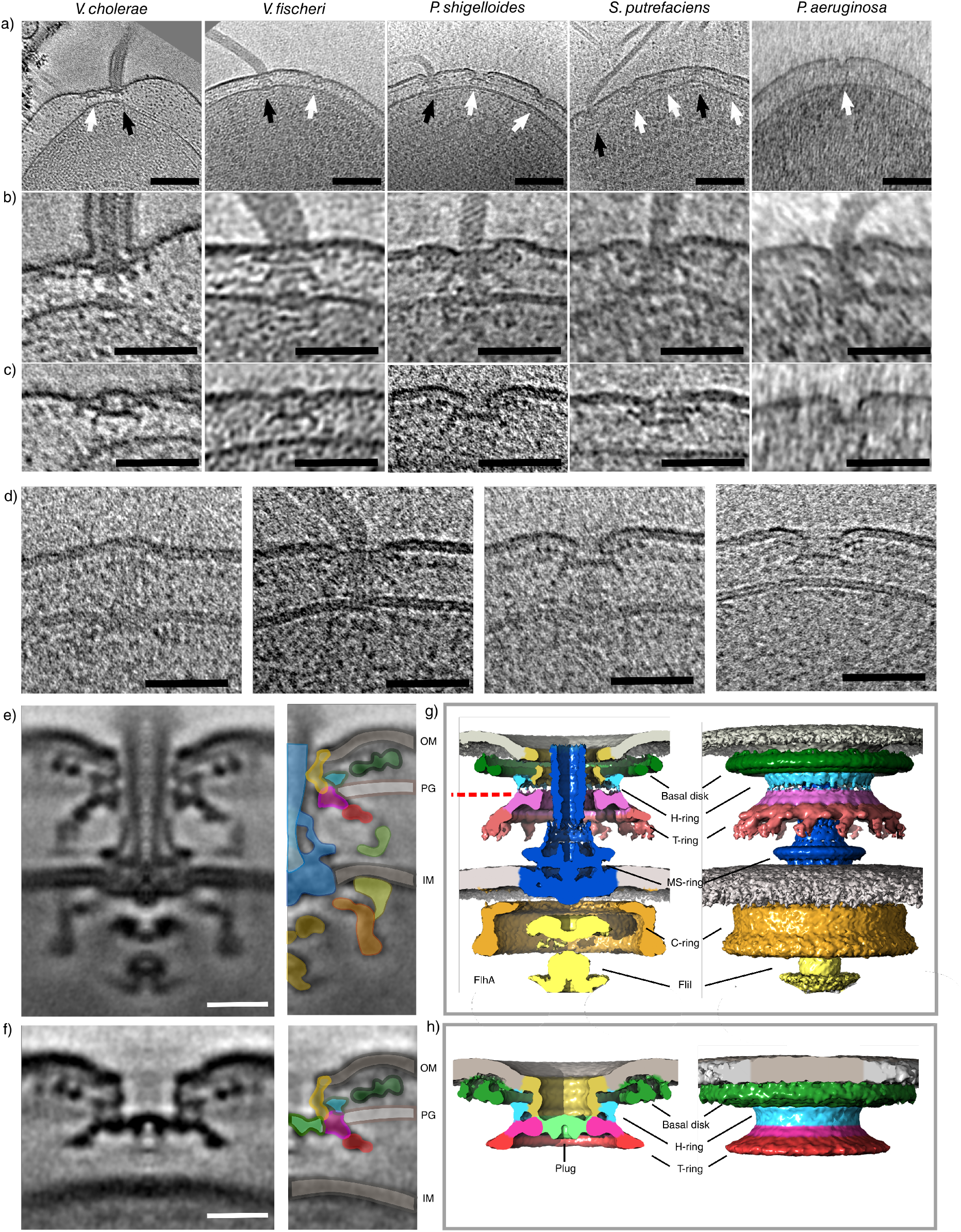
Old γ-proteobacterial cell poles bear the relics of ejected flagella. a) Slices through tomograms showing flagella and relic structures in 5 different γ-proteobacteria, *V. cholera, V. fischeri, P. shigelloides, S. putrefaciens* and *P. aeruginosa*. Black arrows point to full flagella and white arrows point to relics. Scale bar is 100nm. b) Example sub-tomograms of full flagella in the 5 different γ-proteobacteria. Scale bar is 50nm. c) Example sub-tomograms of relics in the 5 different γ-proteobacteria. Scale bar is 50nm. d) Multiple flagellar states are caught in *P. shigelloides* tomograms including precursors that are different from the relic structures. Scale bar is 50nm. e) Central slice through a sub-tomogram average of the *P. shigelloides* polar flagellar motor. Scale bar is 15 nm f) Central slice through a sub-tomogram average of the *P. shigelloides* polar relic. Scale bar is 15 nm g) Isosurface renderings of motor sub-tomogram average shown in e. Dotted red line points to where the plug density sits in the relic. h) Isosurface renderings of relic sub-tomogram average shown in f.

Relics from sheathed and unsheathed γ-proteobacteria had different outer membrane morphologies. Relics from tomograms of sheathed *V. cholerae* and *V. fischeri* were capped by a continuous sealed outer membrane. Ejection of the flagellum in these bacteria must include shearing of the sheath from the outer membrane and subsequent closure of the torn membrane. Evidently the unsheathed species *P. shigelloides, S. putrefaciens*,and *P. aeruginosa* maintain a portal through the outer membrane at all times with their L-ring that is unperturbed upon flagellar ejection, suggesting an as-yet unidentified difference between the L-rings from sheathed and unsheathed bacteria. Occasional relic structures included C-rings, but lacked axial components.

### Multiple assembly precursor states, distinct from relics, are observed in cryo-tomograms

Alongside relics and intact flagella, multiple assembly precursor states were captured in our cryo-tomograms (Fig. 2d). Early precursor structures include fully-formed C-rings and rods but lack P- and L-ring-based outer membrane disks and external structures. However, by far the most common structures where fully formed flagella and relic structures.

### Subtomogram averaging reveals a novel protein density as a putative plug preventing periplasmic leakage

To better understand the relationship of relics to intact motors, subtomogram averaging of both was performed for the *P. shigelloides* structures (Fig. 2e-h). Clear 13-fold symmetry was observed in the stator complexes and MotXY ring of the full motor (Fig. S2), consistent with previous studies (Beeby et al., 2016; Zhu et al., 2017a). The relic structure closely resembled the outer membrane flagellar structure, indicating that the relic is composed of the same proteins as the outer membrane-associated structures from the flagellar motor. In both structures the distance between the MotXY ring (T-ring) and the outer membrane was 19 nm and the diameter of the MotXY ring was 44 nm. In the motor, the diameter of the hole in the outer membrane that the rod exits was 15 nm, the same as the 15 nm hole in the relic outer membrane that is held open by the L-ring despite lack of rod. We conclude that these structures are composed from the same proteins, with the inner membrane components, rod, and hook absent from the relic structure.

Notably, an extra density in the relic structure plugged the P-ring aperture usually occupied by the trans-periplasmic rod. From our observations, this novel plug density appears to be an obligate part of the relic, as no relics were observed lacking a plug, consistent with the need to prevent periplasmic leakage upon filament ejection. Relics from both sheathed and unsheathed bacteria contained the plug protein regardless of the sealing of the outer membrane in the sheathed flagella.

### Both intact flagella and relics follow the same, non-random, placement pattern

We observed that intact flagella and relics appear spaced on a grid at cell poles, suggesting that placement is not random and that both structures share a common grid. The nearest neighbour distances for all intact flagella and relics were determined and found to be on average 64nm, ^∼^20 nm greater than would be expected if distances were constrained by steric clashes of ^∼^45nm-diameter C-rings. To assess whether the placement of both relics and intact flagella is the same, suggesting a common mechanism for placement, histograms were plotted for all distances to nearest neighbours of relics, intact flagella, and random data (figure 3b). The relics and intact flagella shared identical curves, with a much tighter distribution than the randomized dataset, consistent with a common placement mechanism.

**Figure 3:**
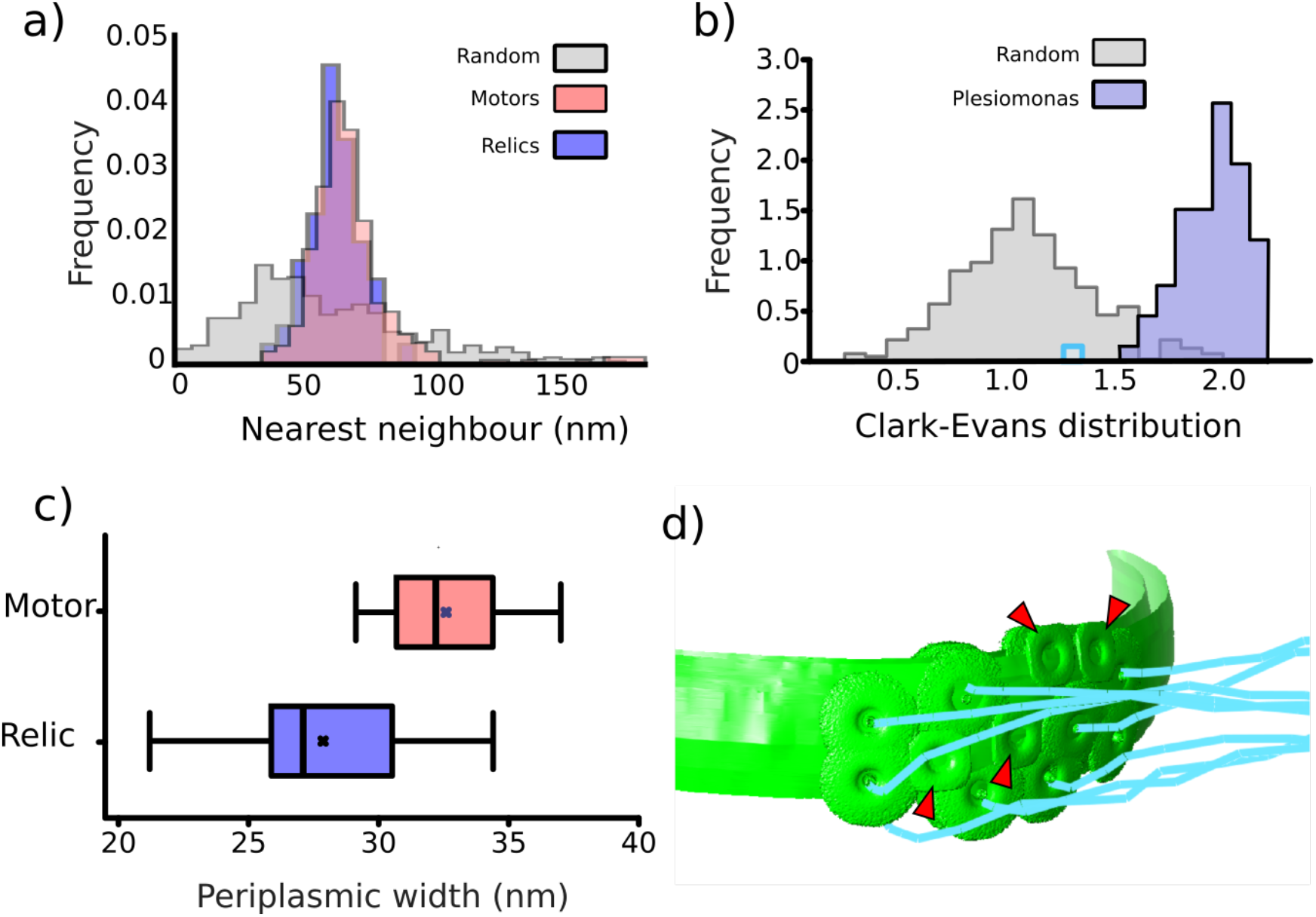
Relic positioning is the same as full flagella indicating a common assembly placement. a) Clarke-Evans distribution of a random synthesized data-set (N=>500) compared to data from tomograms of *P. shigelloides* (N=74). b) Histograms for all nearest neighbours of relics (N=114), motors (N=424) and random synthesized data (N>500). c) Periplasmic distance at position of relics or motors. Box is first and third quartile, the line in the box is the median and the cross is the mean. d) Segmentation of a representative tomogram from *P. shigelloides* showing relative placement of relics and motors. (d) isosurfaces of flagella and relics placed in a membrane model of a representative cell illustrating centrality of relics. Red arrowheads indicate relic structures; blue filaments represent flagella.

To quantify the dispersal of intact flagella and relic structures independent of the number of structures per cell, Clark-Evans distribution analysis (Clark and Evans, 1954) was calculated for each cell pole with more than four intact flagella or relics (N=71) and compared to a synthesised random dataset (figure 3a). Values between 1 and 2.15 suggest a uniform grid while values below 1 suggest clustering. A value close to 1 is a random dispersion. The ^∼^500 synthesised random poles (constrained by the same number of structures and pole area as the real data) had a mean of 1 as expected however, the *Plesiomonas* dataset had a mean close to 2. This suggests that structures at the pole are placed in a non-random distribution in a grid-like arrangement and that the relics and intact flagella are part of the same grid. Peritrichous flagella in *B. subtilis* have previously been shown to be gridded with a Clark-Evans ratio greater than 1 (Guttenplan et al., 2013).

### Relics are no longer tethered to the inner membrane

A common placement mechanism for relics and intact flagella is expected due to their placement within the same grid, likely guided from the cytoplasm. If relic structures are in fact assembly precursors, a connection between the structures and the inner membrane would be anticipated. However, if structures are genuine relics of old intact flagella, flagella placement would have preceeded ejection, and no connection to the inner membrane would be expected to remain.

To determine whether relics are connected to the inner membrane, the distance between the inner membrane and outer membrane disk was measured in intact flagella and relics. The range in periplasmic distances varied in both cases, as was expected from previous work in the *V. alginolyticus* motor (Zhu et al., 2017b). The variation in membrane spacing at the position of relics was far greater than at the motors (13.9nm and 7.9nm range respectively) suggesting no retaining connection between the relic and the inner membrane (figure 3c). As relics are placed in a grid with the intact flagella, this provides evidence that these structures were placed prior to flagellar ejection.

### Ejection is triggered by the depletion of nutrients and not a chemical signal

As ejection is linked to cells at high OD, we confirmed that cells continually diluted with fresh LB, to maintain the optical density below 0.2, retained their flagella. Diluted cells, although in the incubator for the same length of time as undiluted cells that grew to an OD of 1 and had a mean number of 1.7 flagella, still had all flagella (mean of 5.3) (Fig. 4b), confirming that the signal for ejection is a factor of the media of old cells.

**Figure 4:**
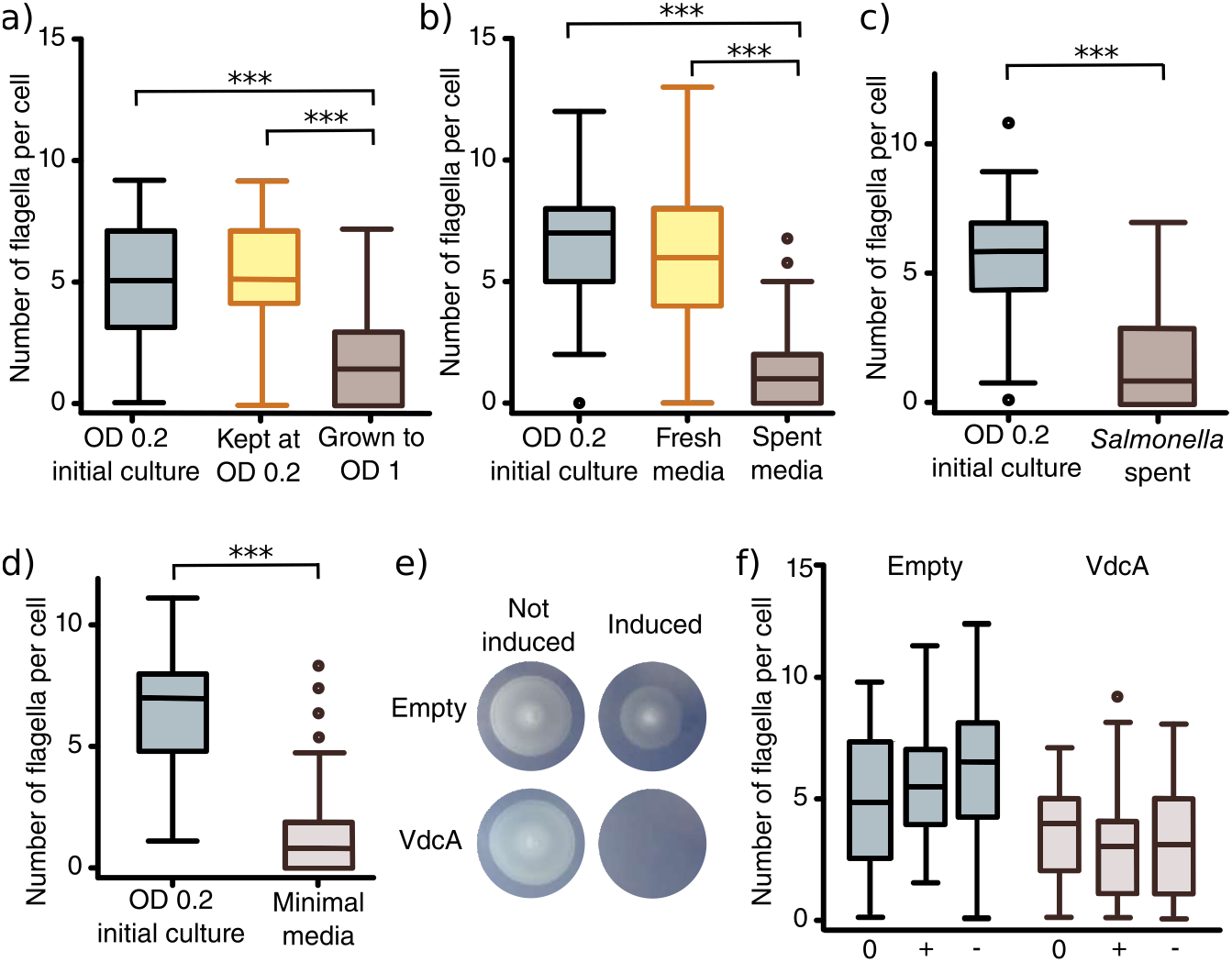
Ejection of the γ-proteobacterial flagellum is triggered by nutrient depletion. a) Cells continually diluted to low cell density do not lose flagella. (*** p < 0.0001, 1-way ANOVA) b) Low cell density cells moved from fresh LB to *P. shigelloides* spent media lose flagella within one doubling time. (*** p < 0.0001, 1-way ANOVA) c) Low cell density cells moved from fresh LB to *S. enterica* spent media lose flagella within one doubling time. (*** p < 0.0001, Unpaired t-test) d) Cells eject flagella when transferred from LB to MOPS-minimal media. (*** p < 0.0001, Unpaired t-test) e) Motility plates showing that cells expressing the cdiGMP maker, VdcA stop swimming. f) Stopping swimming with increased levels of cdG is not due to ejection of flagella. 0= prior to induction, – = not induced, + = Induced. No significant difference was seen between the induced and not induced cells.

To further confirm this, low OD cells were grown in spent media for one doubling time from OD 0.2 to 0.4. Within a single cell-cycle, multiple flagella were lost, with mean number of flagella dropping from 6 to 1. A control culture grown in fresh LB and negative stained at the same optical densities retained all flagella (Fig. 4a). This result suggests that the trigger for flagellar ejection is in the supernatant of cells that have lost flagella but does not differentiate between a signal from a secreted substance, or depletion of nutrients.

To assess whether the signal in the supernatant is a chemical secreted by high OD cells or depletion of a medium component, *P. shigelloides* cells were grown to low OD and transferred to spent supernatant from *Salmonella* grown to stationary phase. Intact flagella were counted after one doubling (Fig. 4c). As with *P. shigelloides* spent media, within one cell division in *Salmonella* spent media, *P. shigelloides* cells had lost multiple flagella (mean number of flagella went from 6 to 2).

To further confirm that the signal is a depletion of nutrients, we moved low OD cells from LB into MOPS minimal media (Fig. 4d). Cells stopped swimming within an hour and within one doubling time the cells had lost most of their flagella as counted by negative stain electron microscopy, with a drop in mean number of flagella from 6 to 1. This suggests that a lack of nutrients is the signal for ejection of flagella and entry into a non-motile state.

### Cyclic di-GMP abolishes motility but does not trigger flagellar ejection

In many bacteria, the transition from motile to sessile states is mediated by concentration changes of the intracellular second messenger cyclic di-GMP (cdG) (Jenal et al., 2017). To determine whether flagellar ejection is mediated by cdG signaling, we overexpressed the cdG-synthesising diguanylate-cyclase, VdcA, from *V. cholerae* in *P. shigelloides* cells and monitored motility and flagellar number. As expected, motility was abolished in *P. shigelloides* cells overexpressing VdcA but not in WT cells harbouring an empty vector and grown under similar conditions (Fig. 4e). Surprisingly, however, when we counted flagella on cells induced to overexpress VdcA we saw no decrease in number of flagella (Fig. 4f). This result demonstrates that flagellar ejection in γ-proteobacteria is not under cdG control.

## Discussion

Here we describe a new mechanism of switching from motile to non-motile states in the polarly flagellelated γ-*proteobacteria*. To achieve this, we used electron microscopy to analyse the structural changes in species that showed characteristic decreases in swimming speeds at high cell density. We found that these bacteria respond to nutrient depletion by shifting from motile to non-motile state by ejecting their flagellar hook and filament into the supernatant. Using ECT we showed that the relics of these ejected flagella are retained by the cell and remain at old cell poles. Sub-tomogram averaging provided a clear view of the relics that were plugged by a novel protein likely to protect against periplasmic leakage. We further showed that plugged relics are found in all of the polarly flagellated γ-proteobacterial species imaged to date: *Plesiomonas shigelloides, Vibrio cholerae, Vibrio fischeri, Shewanella putrefaciens*, and *Pseudomonas aeruginosa*.

### The outer membrane structure is a flagellar relic, not an assembly precursor

Outer membrane “relic” structures could be assembly precursors awaiting insertion of a rod, hook, and filament, and not remnants of old motors. Multiple lines of evidence, however, support the conclusion that these structures are relics of ejected flagella. (1) γ-proteobacterial polar flagella are ejected with correlated decrease in swimming speed, decreased flagella per cell and in the population, and recovery of flagella with hooks from the supernatant. (2) Ejection of flagella at high OD correlates with appearance of greater numbers of relic structures in the outer membrane, and fewer flagella. (3) Localization of relics and flagella at the pole is not random but positioned in a grid-like arrangement. Importantly, the relics are part of the same grid as the flagella. Flagellar placement in multi-polarly flagellated γ-proteobacterial is mediated by FlhF; identical placement pattern of relics and flagella indicate periplasmic relics are positioned by cytoplasmic FlhF, which could only be achieved via the established assembly of a rod around which the (future) relic’s P-ring could form. (4) Furthermore, for the structure to be an assembly precursor, we would expect there to be a connection between the structure and the inner membrane associated with polar localization and assembly. The range in periplasmic distances at the relic however, is far greater than at a full flagellum, indicating absence of a connection from the relic to the inner membrane. (5) In cryo-tomograms, relics are positioned closer to the center of the cell pole than intact flagella (Fig. S1). Presuming FlhF positions flagella as close to the center of the cell pole as possible, this indicates that the structures closer to the center of the pole (i.e., relics) are older than more peripheral structures that are positioned as close as possible to the pole (i.e., newly assembled intact flagella). (6) Appearance of multiple relics at a single pole of *V. cholerae* and *P. aeruginosa*, bacteria that only ever assemble exactly one polar flagellum, is inconsistent with controlled assembly of a single polar flagellum, and best consistent with relics deriving from previously ejected old flagella. The assembly of multiple precursor structures in a stalled state coordinated by as-yet unidentified machinery is less parsimonious with what is known about flagellar assembly. (7) It is established that the P- and L-rings that form the core of the relics require a distal rod for assembly (Kubori et al., 1992); furthermore, polyrods assemble multiple P-rings, showing that the rod, and not the outer membrane, is required for P-/L-ring formation, arguing that the P- and L-rings cannot assembly to form the relic structures alone. Recent work in *V. alginolyticus*, a close relative of *V. fischeri* and *V. cholerae*, suggests that the H-ring is involved in outer membrane penetration in bacteria with sheathed flagella (Liu et al., 2018). (8) Known relic components FlgH, FlgI, FlgO, FlgP, FlgT, MotX and MotY are interspersed with flagellar rod and hook genes in the γ-proteobacterial operon structure (Fig. 5a). The selective production of exclusively these proteins without the many class-II flagellar proteins is highly improbable.

**Figure 5:**
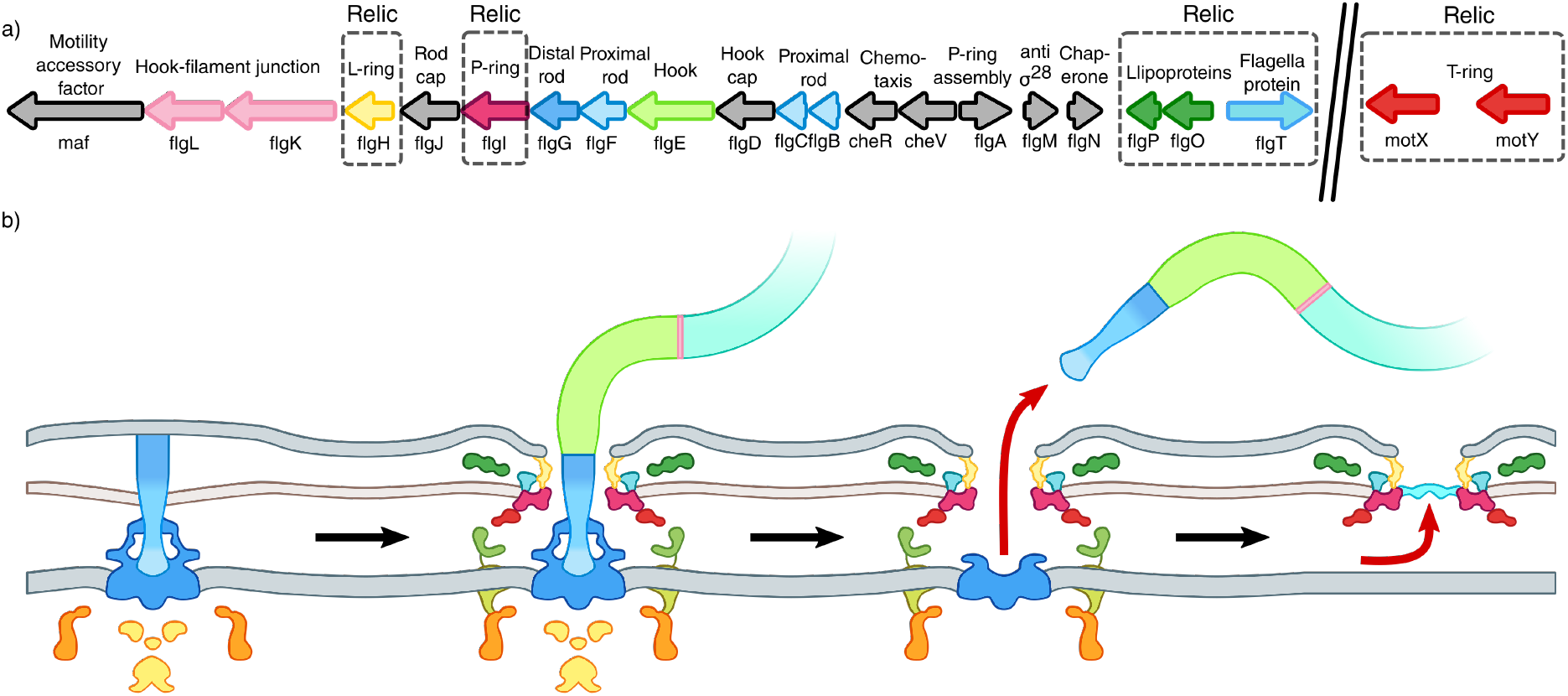
A model for γ-proteobacterial flagellar ejection. a) Operon structure of polar flagellar genes in *P. shigelloides* with dispersed relic genes. Colours match structures in Fig. 2g and h and Fig. 5b. b) Schematic for the assembly and ejection of the γproteobacterial polar flagellum

### Use of a plug protein indicates a specific cellular response

A striking feature of relics is the addition of a “plug” filling the space usually occupied by the distal rod. A targeted response to the ejection of the flagellum in the form of a plug is significant, as this plug may prevent leakage of cellular components. Plug placement is likely rapid, as we observed no relics lacking the plug density. The alternative strategy to prevent periplasmic leakage would be disassembly of the entire relic structure and resealing of the outer membrane, requiring evolution of an intricate machinery for proteolysis of all components. Evolution is a tinkerer (Jacob, 1977), and plugging the hole is a more trivial solution than orchestrated disassembly, particularly given that retaining the relic structures is unlikely to be detrimental to the cell.

What is the identity of the novel plug protein? Using a bioinformatics approach we identified putative plug proteins from a collection of 309 transcripts upregulated at high cell density (Meng et al., 2015; Wagner et al., 2003; Waite et al., 2005). Proteins not conserved in *Vibrio, Pseudomonas, Plesiomonas* and *Shewanella* genera were filtered from the list and the remainder screened for proteins with a SEC signal sequence based on the known SEC secretion of relic proteins FlgH, FlgI, FlgO, FlgP, FlgT, MotX and MotY. This selection yielded four proteins: a putative inner membrane protein, and three of proteins of interest of unknown function upregulated in *Vibrio parahaemolyticus* during early stationary phase: Q87H87, Q87GB3, and Q87PM6 (UniProt identifiers). Bioinformatic inference of function using the STRING database failed to show a specific flagellar association for Q87H87 and Q87GB3, but revealed that Q87PM6 co-occurs across genomes with the T-ring relic component MotY, yet but does not co-occur in operons with MotY, consistent with transcription at a later stage uncoupled from MotY expression. Q87PM6 is co-transcribed with a putative anti-σ^F^ factor antagonist, capsule transport protein OtnA, and a putative capsular polysaccharide biosynthesis protein. Potential homologues of Q87PM6 identified by PSI-BLAST align to a common OmpA-like domain on the C-terminus of the protein consistent with peptidoglycan association. In addition, Q87PM6 was experimentally shown to interact with putative OtnA and TolR. We speculate, therefore, that Q87PM6 forms part of the γ-proteobacteria relic plug, and will test this hypothesis in future studies.

We expect that in the relatively well-studied case of *Caulobacter crecentus* flagellar ejection a similar plugging mechanism may occur (Jenal 2000). Although an active mechanism for rearrangement of the outer membrane after flagella ejection is likely to exist to prevent leakage of periplasmic components out of the cell, at present it is unclear what this might be and what the fate of the outer membrane components is after ejection. The *C. crecentus* flagellar motor, however, does not contain large outer membrane disks (Chen et al., 2011) and so the visualization of a *Caulobacter* relic, made up of only the P and L-rings would prove difficult with ECT.

### Mechanism of flagellar ejection

Our study does not identify the mechanism of flagellar ejection. Nevertheless, the results indicate that the trigger for ejection is nutrient depletion, which, together with identification of a plug protein, argues against the mechanism being mechanical breakage at the base of the hook. Indeed, we routinely monitor γ-proteobacterial motility in more viscous media, which would put more strain on the motor, yet do not observe abolished swimming motility resulting in viscosity changes. In *C. crescentus*, the protease ClpAP is associated with degradation of FliF at the time of flagellar ejection, although it is unclear whether this is causal, or caused by, ejection (Ardissone and Viollier, 2015). To identify proteins involved in the γ-proteobacterial ejection process a test will be to identify ejection suppressors that retain the ability to swim upon transfer to minimal medium. These suppressor mutations will enable identification of proteins directly involved in ejection, or signaling components.

### The selective benefit of flagellar ejection

The γ-proteobacteria with sodium-driven motors eject their flagella and enter a non-motile state, unlike *Salmonella*. As ejection occurs in nutrient limited environments this may be a mechanism for bacteria cultures to conserve energy by cutting back on its more extravagant energy expenditures such as flagellar motility.

Why polarly flagellated γ-proteobacteria eject motors under nutrient depletion, whereas *Salmonella* and *E. coli* simply cease assembly, leading to a decrease in flagellar number over time due to dilution, is unclear, yet being peritrichous, *Salmonella* and *E. coli* also lose flagella at the onset of stationary phase by dilution as the cell elongates. With the polar flagellates, however, motors are polar, and therefore not diluted as the cell elongates. Rather, one daughter cell inherits an entire set of flagella. Ejection, therefore, is a drastic but potentially necessary survival mechanisms for those daughter cells that inherit old, already-flagellated poles. *V. fischeri* has previously been shown to lose flagella upon colonisation of the squid light cavity and regrow them when exposed to fresh seawater (Ruby and Asato, 1993), although how this observation generalizes over other γ-proteobacteria is unclear. Flagellation levels have been shown to decrease as bacteria enter stationary phase. We suggest that flagellar ejection is a mechanism to reduce the number of flagella on a cell pole. Not only do active flagella require energy for torque generation, but flagellar assembly is a constant process and a constant drain on cell energy (Paradis et al., 2017). Though drastic, decreasing the number of flagella by ejection would be an effective energy conservation measure.

### Model for assembly and ejection of the sodium-driven polar flagellum

Our results enable interpretation of the multiple states of the flagellum caught in tomograms (Fig. 2d) we propose a model for the steps of assembly and ejection of the polar γ-proteobacterial flagellum (Fig. 5b). After assembly of the MS-ring, C-ring and T3SS, the rod is assembled. Recent work in *Salmonella* showed that polymerization of the rod is stopped by hitting into the outer membrane (Cohen et al., 2017). In sub-tomograms of this stage, the rod appears to be hitting the outer membrane perpendicular and pushing on it forming a bulge in the outer membrane. We believe that the same mechanism for stopping rod polymerisation in the enteric motors is found in polar flagellar motors. Once the P- and L- rings form around the rod, FlgO, FlgP, FlgT, MotX and MotY can assemble on this platform, and the hook and filament can be assembled through the channel in the outer membrane formed by the L-ring. Upon nutrient depletion, however, the filament, hook and distal rod are released and a plug protein fills in the leftover gap. The inner membrane and C-ring components are finally cleared away. The ejection of flagella in nutrient depleted environments and subsequent rebuilding when exposed to fresh nutrients is a novel mechanism for the transition between a planktonic and a sessile state shown here to be widespread amongst the sodium-driven polar flagellated γ-proteobacteria.

## Methods

### Cell Growth

*Plesiomonas shigelloides* was grown in LB at 37°C, 200RPM overnight in 14ml aeration tubes. The next day, cultures were diluted 1:100 for experiments. Cells were grown to an OD of 0.2 unless specified.

*Shewanella putrefaciens* CN-32 (*ΔflhG*) was grown in LB at 30°C, 200RPM overnight in 14ml aeration tubes. The next day, cultures were diluted 1:100 for experiments. Cells were grown to an OD of 0.5.

*Vibrio fischeri* was grown in LBS at 30 degrees, 200RPM overnight in 14ml aeration tubes. The next day, cultures were diluted 1:100 for experiments. Cells were grown to an OD of 0.3.

*Vibrio cholerae* N16961 was grown in LB for 18h, harvested by centrifugation and washed twice with sterile filtered and autoclaved environmental sea water before introduced into a 50ml sea water microcosm at an OD_600_=2.5. Microcosms were kept at low agitation at 4°C for one month prior to imaging.

*Pseudomonas aeruginosa* PAOI persister cells were obtained from overnight stationary phase culture treated with Carbonyl cyanide *m*-chlorophenylhydrazone (CCCP) 200 μg/mL for 3h in LB, followed by exposure to Ciprofloxacin 5 μg/mL for 3h in sodium-phosphate buffer (10 mM, pH 7.4) as described previously (Grassi et al., 2017).

| Strain or Plasmid | Source |
| --- | --- |
| *Plesiomonas shigelloides* WT | DSMZ |
| *Shewanella putrefaciens* CN-32 (ΔflhG) | (Schuhmacher et al., 2015) |
| Vibrio fischeri WT | DSMZ |
| Vibrio cholerae N16961 WT | Simon Ringgaard (MPI Marburg) |
| Pseudomonas aeruginosa PAOI WT | Kylie Watts (Loma Linda University) |
| E.coli WM3064 | W. Metcalf, (University of Illinois) |
| pBTOK | (Rossmann et al., 2015a) |
| pBTOK-VdcA | This study |

### Swimming Speed Videos

To measure bacterial swimming speeds at different cell densities, cells were grown as stated above and diluted to an OD_600_ of 0.2 immediately prior to imaging. Cells were imaged at 100x oil immersion on a MT4000 series light microscope (Meiji Techno) with a DMK23G618 digital camera connected to IC Capture software. Bacterial cells were tracked for 10 seconds using Fiji and converted to swimming velocities using custom scripts (Schindelin et al., 2012). Three videos were taken at each time point providing over 1000 traces and this was done in triplicate. Error bars indicate standard error.

### Negative stain EM

3µl of cells as described above, were placed on glow-discharged continuous-carbon grids (Taab Laboratory Equipment Ltd.). The grids were stained with 2% Uranyl Acetate and stored at room temperature prior to imaging. Negative stain images were collected on either an FEI Tecnai G2 Spirit BioTWIN (Tungsten) using an Eagle CCD camera or a Technai T12 TWIN (LAB6) using a TVIPS F216 CCD camera. Images were viewed using 3DMOD and number of flagella per cell or number of hooks was counted manually.

### PEG precipitation to recover ejected flagella

Cells grown overnight were spun down three times for 30 minutes at 4000RPM and supernatant was carefully decanted. The supernatant was assessed with light microscopy to make sure no cells remained. 2% PEG 20,000 was added to the supernatant and stirred at RT for >1hour. Flagella were recovered by centrifugation at 4000RPM for 20 minutes and dialysed to remove PEG.

### SDS-gel electrophoresis of recovered flagella

After PEG precipitation, flagella were boiled in LDS and run on a 4–12% Bis-Tris gel (Thermo-Fisher, Hillsboro, OR, USA) and stained with InstantBlue (Sigma-Aldrich, Taufkirchen, Germany).

### Mean number of flagella in a population

CFU at each OD was calculated by plating cells on LB or LBS agar and counting the number of colonies. The mean number of flagella at each time point, from negative stain images was multiplied by the total number of cells in the population.

For all experiments at least three biological replicates with 30–50 cells per time point in each were imaged.

### Electron Cryo-Tomography Grid Preparation

For *P. shigelloides, S. putrifaciens and V. fischeri*: Cells were pelleted by centrifugation at 6000RPM for 5 minutes and resuspended to an OD600 of 11. Cells were mixed with gold fiducials coated in BSA and 3µl of the cell fiducial mixture was applied to freshly glow discharged R2/2, 300mesh gold Ultrafoil or copper Quantifoil R2/2 grids (Quantifoil Micro Tools GmbH, Germany). A Vitrobot Mark IV was used to freeze grids in an ethane/propane cryogen at 100% humidity.

*V. cholerae* and *P. aeruginosa* cells from cultures were directly mixed with protein A-treated 10 nm colloidal gold solution (Celll Microscopy Core, Utrecht University, Utrecht, The Netherlands). After mixing, aliquots of 3 µl were applied to freshly plasma-cleaned R2/2, 200mesh copper Quantifoil grids (Quantifoil Micro Tools GmbH, Germany). Plunge freezing was carried out in liquid ethane using a Leica EMGP (Leica microsystems, Wetzlar, Germany). During 1s blotting time, the blotting chamber was set at room temperature (20°C) with 95% humidity.

### ECT Data Acquisition

Exploratory tomograms of *Plesiomonas, Shewanella* and *Vibrio fischeri* were collected on an FEI F20 with a Falcon II detector (Thermo Fisher Scientific (formerly FEI), Hillsboro, OR, USA). A total fluence of 120e-/ Å^2^ was used and a defocus of –4µm. Tomograms were collected at a tilt range of +-53 degrees with 3° tilt increments and a pixel size of 8.28 Å.

Exploratory tomograms of *V. cholerae* and *P. aeruginosa* was performed on a Titan Krios transmission electron microscope (Thermo Fisher Scientific (formerly FEI), Hillsboro, OR, USA) operating at 300 kV. Tomograms were recorded with a Gatan K2 Summit direct electron detector (Gatan, Pleasanton, CA) equipped with a GIF-quantum energy filter (Gatan) operating with a slit width of 20eV. Images of *V. cholerae* and *P. aeruginosa* were taken at a nominal magnification of 42,000x and 26,000x respectively, which corresponded to a pixel size of 3.513 Å and 5.442 Å. Using UCSFtomo software, all tilt series were collected using a bidirectional tilt scheme of +-53 degrees (*V. cholerae*) and of +-60 degrees (*P. aeruginosa*), respectively. Defocus was set to –8 µm.

The *P. shigelloides* data-set for subtomogram averages were collected at the Francis Crick Institute on a Titan Krios operating at 300 kV with a post-column GIF using a 20eV slit and a Gatan K2 summit (Gatan, Pleasanton, CA) collecting 4 frames per exposure. A pixel size of 2.713 Å with a total fluence of 54e/A2 and a defocus range between −3 and −5.5µm was used. The bi-directional tilt series was collected using FEI’s Tomo4 (Thermo Fisher Scientific (formerly FEI), Hillsboro, OR, USA) with a tilt range of +-53 degrees and a tilt-increment of 3 degrees.

### Tomogram reconstruction

For the *P. shigelloides* dataset, Frames were aligned using UnBlur (Grant and Grigorieff, 2015). Tomoctf (Fernández et al., 2006) was used to estimate CTF-parameters (tomoctffind) and correct stacks (ctfcorrectstack). Fiducials were picked from CTF-corrected stacks and tracked in IMOD (Kremer et al., 1996; Mastronarde, 1997). The aligned, CTF-corrected stacks and fiducial models were used with Tomo3d to reconstruct using SIRT (Agulleiro and Fernandez, 2011, 2015).

For *V. fischeri* and *S. putrificiens* Tomograms were reconstructed using RAPTOR (Amat et al., 2008).

For V. cholera and P. aeruginosa, drift correction, fiducial-tracking based tilt series alignment of tomograms were done using software package IMOD(Kremer et al., 1996; Mastronarde, 1997). Tomograms were reconstructed using SIRT with iteration number set to 4.

### Subtomogram Averaging

Subtomograms were picked manually. Template-free alignment was carried out in PEET by superimposing all manually picked subtomograms allowing no shifts or rotations for an initial reference. Initial alignment and averaging was achieved using binned subtomograms. Unbinned subtomograms were used for further refinement. As the rod length varied in the full motor, sub-tomogram averaging was carried out twice with custom masks applied to either the top ring structures or the cytoplasmic structures. As clear 13-fold symmetry was observed in the full motor (fig. S2), 13-fold rotational averaging was applied using custom scripts. The two individual averages for the top and bottom segments were then merged into a composite structure as in (Beeby et al., 2016). For comparisons, 13-fold rotational averaging was applied to the relic structure. FSC curves were determined from the final, unsymmetrised averages using PEET’s calcFSC (fig. S2).

### Segmentation

Semi-automatic segmentation of the membranes was performed in IMOD using the Sculpt drawing tool followed by the linear interpolator.

### Periplasmic distance measurements

Contours were drawn between the inner membrane and the first ring of the outer membrane disks using 3dMOD (IMOD package) and distances were measured.

### Clark-Evans distribution analysis

Clark-Evans ratios were calculated per cell with more than four flagella (N=71) (Clark and Evans, 1954). To calculate the Clark-Evans distribution, the mean nearest neighbour distance was divided by half the square root of the mean cell pole density. The mean nearest neighbour distance was calculated using custom scripts and the cell pole area density was calculated for each pole. The area of each pole was calculated by adding the diameter of one C-ring (45nm) to the maximum X and Y coordinates of any motor or relic structure and these coordinates were used to calculate the area of an ellipse. To calculate the mean cell pole density, each pole area was divided by the number of structures at that pole.

### Nutrient depletion assay

Cells grown overnight were removed from the supernatant by centrifugation. The supernatant was subsequently filtered using a 0.45µm filter. Plesiomonas cells grown to low OD_600_ (0.2) were spun down and resuspended in the filtered, spent medium. Negative stain grids were made prior to resuspension and after one doubling time in spent media.

### Minimal media assay

*Plesiomonas* cells grown to low OD (OD 0.2) were spun down and resuspended in MOPS-minimal medium (40mM MOPS pH 7.4, 4mM Tricine, 0.1mM FeSO_4_·7H_2_O, 9.5mM NH_4_Cl, 0.28 mM KCl, 0.53 mM MgCl_2_·6H_2_O, 50 mM NaCl, 10mM KH_2_PO_4_, 0.005% MgSO_4_, 0.0005% MnCl_2_·4H_2_O, 0.0005% FeCl_3_, 0.5% glucose). Cells were examined by light microscopy after one hour to see if they were swimming. Negative stain grids were made prior to resuspension and after one doubling time in spent media.

### Strain construction

Inducible expression of proteins in *Plesiomonas shigelloides* was carried out using the pBTOK vector (Rossmann et al., 2015b). DNA manipulations were carried out using appropriate kits (VWR International GmbH, Darmstadt, Germany) and enzymes (Thermo Fisher, St Leon-Rot, Germany). The oligonucleotide pair CGC TCT AGA AGG AGG GCA AAT GTG ATG ACA ACT GAA GAT TTC AAA A & TCC GGG CCC TTA GAG CGG CAT GAC TCG ATT G (Sigma-Aldrich, Taufkirchen, Germany) were used to amplify the genes encoding the diguanylate cyclase VdcA (VCA 0956) from *V.cholerae*. Subsequently, the plasmid was constructed by restriction digest of the vector and the inserts with XbaI and PspOMI and subsequent ligation with Taq Ligase (Thermo Fisher, St. Leon-Rot, Germany).

Plasmids were transformed into chemically competent *E.coli* strains. Subsequent delivery to *Plesiomonas shigelloides* occurred by conjugation from E. coli WM3064.

### Motility plates

*P.shigelloides* cells harbouring either empty pBTOK vector or expressing the diguanylate cyclase VdcA were stabbed into motility plates (10g/L Bacto-Tryptone, 5g/L NaCl, 3g/L Bacto-Agar) containing 25µg/ml Kanamycin and either no inducer or 2ng/ml Anhydrotetracycline and grown at 37°C for 6 hours.

### cdiGMP over-expression

*P.shigelloides* cells harbouring either empty pBTOK vector or expressing the diguanylate cyclase VdcA were grown with 25µg/ml Kanamycin to OD_600_ 0.2. Cells were then split in two and either not induced or induced with 2ng/ml Anhydrotetracycline until an OD_600_ of 0.4 was reached. Negative stain grids were made prior to and after induction.

### Bioinformatic analysis for plug candidates

Initially, 309 protein sequences were identified from the literature that were upregulated during late early stationary phase in *Vibrio parahaemolyticus (Meng et al., 2015)*, upregulated in confluent biofilms in *Pseudomonas aeruginosa (Waite et al., 2005)*, and upregulated quorum sensing regulated transcripts in early stationary phase in *Pseudomonas aeruginosa (Wagner et al., 2003)*.

From tomography data, it is clear that the plug must be exported into the periplasm, and must therefore contain a signal sequence. The sequences were analysed with SignalP 4.0 (Petersen et al., 2011) using the gram negative organism group to predict the probability of a signal sequence being present. We filtered out any sequences without a signal sequence, which was defined as having a S-score below the 0.5 threshold along all parts of the sequence. 52 candidates were identified at this stage.

A PSI-BLAST (Altschul et al., 1997) search was performed on the 52 sequences across the non-redundant protein sequence databases of the *Vibrio, Pseudomonas, Plesiomonas* and *Shewanella* genera for which relics have been observed. The default BLOSUM62 matrix with 11 gap penalty and 1 gap extension was used as the initial scoring matrix. The maximum target sequences were set to its maximum value of 20,000 to ensure that all possible distant homologues could be identified. From the output, we filtered out any sequences where there was no homologue in any of the genera of significant E-value (0.005). 23 candidates were identified at this stage (Supplementary Table 1).

2 out of the 23 protein sequences were uncharacterised in all 4 genera. However, in order to account for the possibility of the plug protein having dual functionality, we inputted the remaining 23 sequences into STRING (Szklarczyk et al., 2017) search to identify interacting partners or co-expressed proteins that are involved in flagella regulation to verify the identity of potential plugs.

## Acknowledgements

This work was funded by a Medical Research Council grant MR/P019374/1 to MB, Medical Research Council Ph.D. Doctoral Training Partnership award grant number MR/K501281/1 to JLF, a research fellowship of the German Research Foundation (DFG project number 385257318) to FMR, TH831/5–1 to KMT, Netherlands Organisation for Scientific Research (NWO) BBOL.16.007 to AB, German Academy of Sciences Leopoldina (Fellowship Programme LPDS 2017–01) to SB, P.B.R. is supported by the Francis Crick Institute, which receives its core funding from Cancer Research UK (FC001143), the UK Medical Research Council (FC001143), and the Wellcome Trust (FC001143). The authors would like to thank Bonnie Basler, Ned Ruby, Kelly Hughes, Teige Matthews-Palmer, Louie Henderson, Eli Cohen for insightful discussions, and Paul Simpson for electron microscopy assistance. We thank Christoph Diebolder for image acquisition at the Netherlands Centre for Electron Nanoscopy in Leiden (NeCEN) and helpful advice.

